# Early candidate biomarkers found from urine of astrocytoma rat before changes in MRI

**DOI:** 10.1101/117333

**Authors:** Yanying Ni, Fanshuang Zhang, Manxia An, Wei Yin, Youhe Gao

**Author notes:** Corresponding author: Youhe Gao, Department of Biochemistry and Molecular Biology, Beijing Normal University, Gene Engineering and Biotechnology Beijing Key Laboratory, Beijing, 100875, PR. of China.

## Abstract

Astrocytoma is the most common aggressive glioma and its early diagnosis remains difficult. Biomarkers are changes associated with the disease. Urine, which is not regulated by homeostatic mechanisms, accumulates changes and therefore is a better source for biomarker discovery. In this study, C6 cells were injected into Wistar rats brain as astrocytoma model. Urine samples were collected at day 2, day 6, day 10 and day 13 after injection, and the urinary proteomes were analyzed. On the 10th day, lesions appeared in magnetic resonance imaging. On the 13th day, clinical symptoms started. But differential urinary proteins were changed with the development of the astrocytoma, and can provide clues even on the 2nd and 6th day. Twenty-seven differential proteins with human orthologs had been reported to associate with astrocytoma. Thirty-nine proteins were verified in four more rats as candidate biomarkers of astrocytoma using multiple-reaction monitoring. A panel of differential urinary proteins may provide early biomarkers for diagnose of astrocytoma.

## Introduction

Gliomas are the most frequent primary tumors of the central nervous system, accounting for more than 60% of all brain tumors [1]. Among them, glioblastoma (GBM) is the most common aggressive glioma, and the average survival rate remains low [2]. The standard combination of surgery and radiotherapy or chemotherapy can only extend the progression-free survival time for 12–18 months, with a high risk of recurrence [3]. In addition, the incidence of astrocytoma has increased annually by 1–2% in the past years [4]. Therefore, a precise early diagnosis of astrocytoma is crucial for the early determination of appropriate therapeutic strategies.

Changes associated with physiological or pathophysiological processes are the most fundamental characteristic of biomarkers [5]. Urine which is not regulated by homeostatic mechanisms, has the potential to accumulate changes and serve as a better source for disease biomarker discovery [6]. However, urine can be affected by various factors, such as age, diet, and drugs [7, 8]. To minimize these confounding factors, animal models were used to mimic the pathophysiological changes of diseases to search for valuable clues, instead of clinical samples [9].

The rapidly proliferating rat C6 cell line is morphologically similar to GBM when injected into the brain of neonatal rats [10]. The C6 model is closer to the usual histological characteristics of spontaneous GBM when using Wistar rats rather than other strains, including nuclear pleomorphism, high mitotic index, hemorrhage, necrosis and parenchymal invasion [11]. In this study, urine samples were used to identify the candidate biomarkers involved in GBM diagnosis, and further monitor prognosis. The urinary proteome at 2 days, 6 days, 10 days and 13 days was analyzed. This study will not only benefit patients with GBM but also give us a more comprehensive understanding of the clinical significance of urine (Fig. 1).

**Fig 1.**
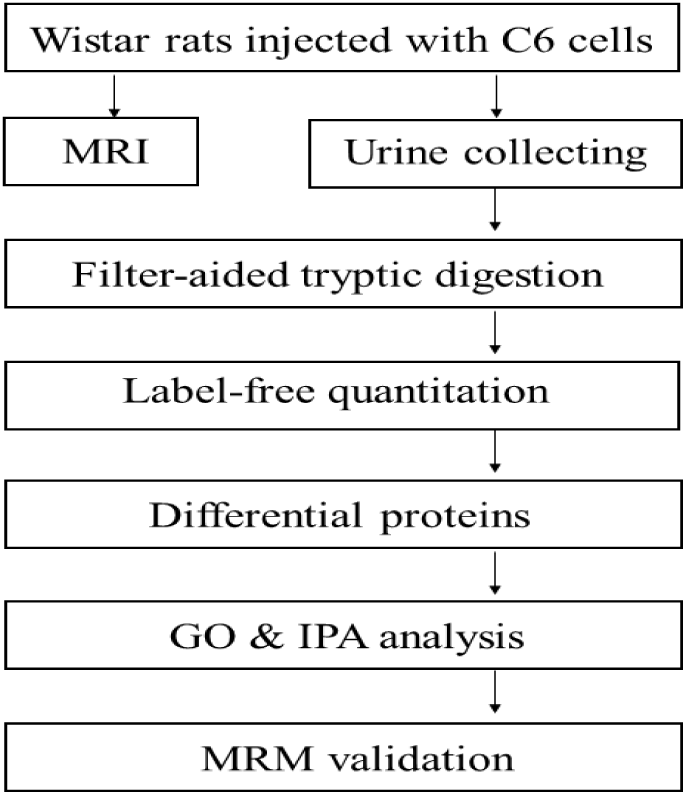
Workflow of protein identification in the GBM rats model. Urine samples were collected from the control and GBM groups. The proteins were analyzed by label-free proteomic analysis. The differential proteins were analyzed by GO and IPA. MRM was used to validate the key differentially expressed proteins.

## Methods

### Ethics statement

All protocols involving animals were approved by the Institute of Basic Medical Sciences Animal Ethics Committee, Peking Union Medical College. Male Wistar rats weighing 200 to 230 g were obtained from the Institute of Laboratory Animal Science, Chinese Academy of Medical Science & Peking Union Medical College. All animals were housed with free access to water and a standard laboratory diet under controlled room temperature (22±1°C) and humidity conditions (65-70%). All efforts were made to minimize suffering.

### Cell culture

Rat C6 cells were obtained from the Chinese Academy of Medical Science & Peking Union Medical College (Beijing, China) and cultured at 37°C in DMEM, supplemented with 10% fetal bovine serum, 100 IU/mL penicillin and 100 μg/mL streptomycin in a humidified atmosphere of 5% CO2 air (Thermo Fisher Scientific, Inc., Waltham, MA, USA) [12]. Cells in the logarithmic growth phase were used for the experiment.

### Rat C6 astrocytoma model

Male Wistar rats were randomly divided into two groups (20 rats in each group). The astrocytoma model was established as described previously [13]. Briefly, under general anesthesia with 2% pelltobarbitalum natricum (40 mg/kg body weight), the skull was drilled at 1 mm anterior to the anterior fontanelle and 3 mm right of the midline, and 10 μl of cell suspension containing 106 cells was injected using a microsyringe (SHANGHAI HIGH PIGEON INDUSTRY TRADE & Co., LTD CHINA). Rats in the control group were administrated PBS only. The rats’ weights were monitored twice a day. Rats with similar clinical symptom were chosen for further analysis, which will help to find biomarkers in the same period of GBM.

### Magnetic resonance imaging analysis

MRI was performed using a 7 T MRI animal system (Agilent, US) on days 6, 10 and 13 after tumor cell implantation. Rats were monitored under anesthesia. The parameters were set as follows: T1 (TR/TE = 500/15.69 ms, 16 slices with FOV 40 × 40 mm^2^, matrix = 256 × 256). Acquisition duration was 3 min and 44 s; T2-weighted sequence (TR/TE =3500 ms/72.00 ms, with repetition time 1,000 ms, matrix = 256 × 256) [14].

### Brain histopathology

Rats were sacrificed using overdose anesthesia and perfused with 4% paraformaldehyde via the left ventricle. After perfusion with 0.9% saline and ice-cold 4% paraformaldehyde, the brains were removed and post-fixed in 4% paraformaldehyde in PBS overnight, and then the tissues were embedded in paraffin, sliced at 2−3 μm, and stained with hematoxylin and eosin (H&E) [15].

### Urinary protein sample preparation and LC-MS/MS analysis

Proteins were extracted as follows [16]: (1) centrifugation at 2000×g and 12000×g for 30 minutes at 4°C, respectively; (2) precipitation by ethanol overnight at 4°C; (3) dissolution by lysis buffer (8 M urea, 2 M thiourea, 25 mM dithiothreitol and 50 mM Tris).

For each urine sample, some proteins were separated using sodium dodecyl sulfate polyacrylamide gel electrophoresis (SDS-PAGE), while some proteins were digested by trypsin (Trypsin Gold, Mass Spec Grade, Promega, Fitchburg, WI, USA) in 10-kD filter units (Pall, Port Washington, NY, USA) [17]. UA (8 M urea in 0.1 M Tris-HCl, pH 8.5) was added to wash the proteins at 14000×g for 20 min at 18°C. NH4HCO3 was added subsequently to wash the protein. Then, the urinary proteins were denatured by dithiothreitol and alkylated by iodoacetamide. The proteins were digested with trypsin at 37°C for 14 hours. The collected peptide mixtures were desalted using Oasis HLB cartridges (Waters, Milford, MA) and then dried by vacuum evaporation (Thermo Fisher Scientific, Bremen, Germany).

The digested peptides were acidified with 0.1% formic acid, then loaded onto a reversed-phase micro-capillary column using a Waters UPLC system. The MS data were acquired using Thermo Orbitrap Fusion Lumos (Thermo Fisher Scientific, Bremen, Germany). Animals (n=3) with the same clinical manifestations were randomly chosen from the astrocytoma group. Each sample was analyzed three times.

### Protein identification and label-free quantitation

The MS/MS data were analyzed using Progenesis and Masccot (version 2.4.1, Matrix Science, London, UK). The parameters were set as follows: Swiss-Prot rat database (551,193 sequences; 196,822,649 residues); the fragment tolerance was 0.05 Da; the parent ion tolerance was 0.05 Da; the precursor mass tolerance was 10 ppm; two missed trypsin cleavage sites were allowed; and peptide identifications containing at least 2 identified peptides were accepted, as in a previous study [18]. Carbamidomethylation of cysteines was set as a fixed modification, and oxidation of methionine and protein N-terminal acetylation were set as variable modifications. After normalization, the fold change of abundance was used to analyze differential proteins between the control group and the astrocytoma group [19, 20].

### Multiple-Reaction Monitoring analysis

Some differential proteins were validation using multiple-reaction monitoring (MRM). For targeted proteomic analysis, the dat file of GBM urine generated by conventional LC/MS/MS were imported into Skyline version 1.1. software. Skyline was employed to select the most intense peptide transitions, up to four or five transitions per peptide were selected and were traced on a QTRAP 6500 mass spectrometer (AB Sciex, Massachusetts, USA). All MS data were imported into Skyline, which was used for further peptide transitions selection and abundance calculations.

### Gene Ontology and Ingenuity Pathway Analysis

All differentially expressed proteins identified between the control and astrocytoma groups were assigned a gene symbol using the Panther database (http://www.pantherdb.org/) and analyzed by Gene Ontology (GO) based on the molecular function, biological processes, and cellular component categories. For biological processes, all differentially expressed proteins were also analyzed by the IPA software (Ingenuity Systems, Mountain View, CA, USA) for pathway analysis, and network analysis.

### Statistical analysis

Clinical symptoms, proteome analysis and MRM data were analyzed using the SPSS16.0 software package for statistical analysis. Comparisons between independent groups were conducted using one-way ANOVA followed by post hoc analysis with the least significant difference (LSD) test or Dunnett’s T3 test. P-values of less than 0.05 were considered different.

## Results and Discussion

### Clinical symptoms

After injection with C6 cells, there was no difference in clinical manifestations. However, on the 13th day, some GBM rats showed twitching, tachypnea, drowsiness and low symptom scores. The rats in the control group did not show any symptoms (Fig. 2A). All rats gained weight steadily after tumor cell implantation. Compared with control rats, the GBM rats’ weights were significantly lower on the 13th day, and the weight was 1.04-fold higher in the control group than in the GBM group (Fig. 2B).

**Fig 2.**
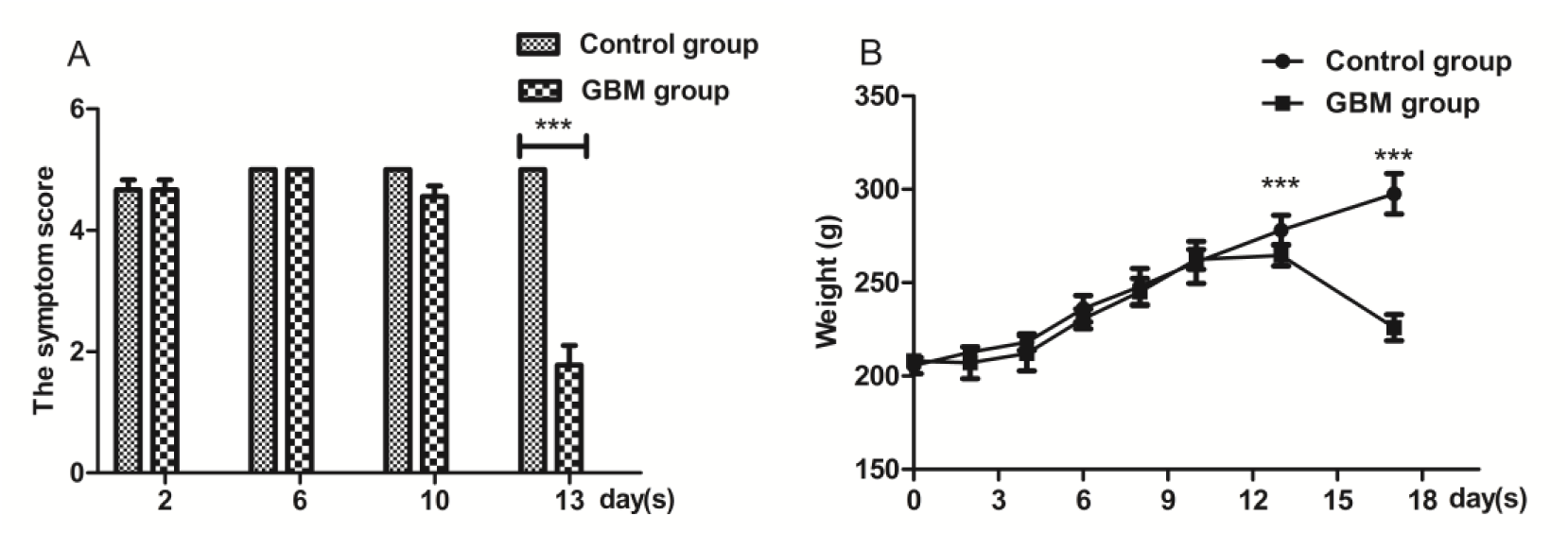
Clinical parameters of rats in control and GBM rats. (a) The symptom scores of GBM rats were significantly decreased on 13th day after tumor cells injection (P=0.001). (b) The weight of GBM rats were significantly decreased on 17th day (P<0.001).

### MRI analysis

Images of rat brains on the 6th day, 10th day and 13th day were obtained. For MRI imaging, the rats were in a supine position. Compared to the left brain, the right brain injected with tumor cells showed no obvious lesion on the 6th day after tumor cell implantation (Fig. 3A). On the 10th day, there was strong enhancement with an obscure boundary in the right brain on the T2 images (Fig. 3B). On the 13th day, the strong enhancement lesions were largger, and the midline shifted on the T2 images, showing an obvious effect of an intracranial placeholder, such as a midline shift (Fig. 3C).

**Fig 3.**
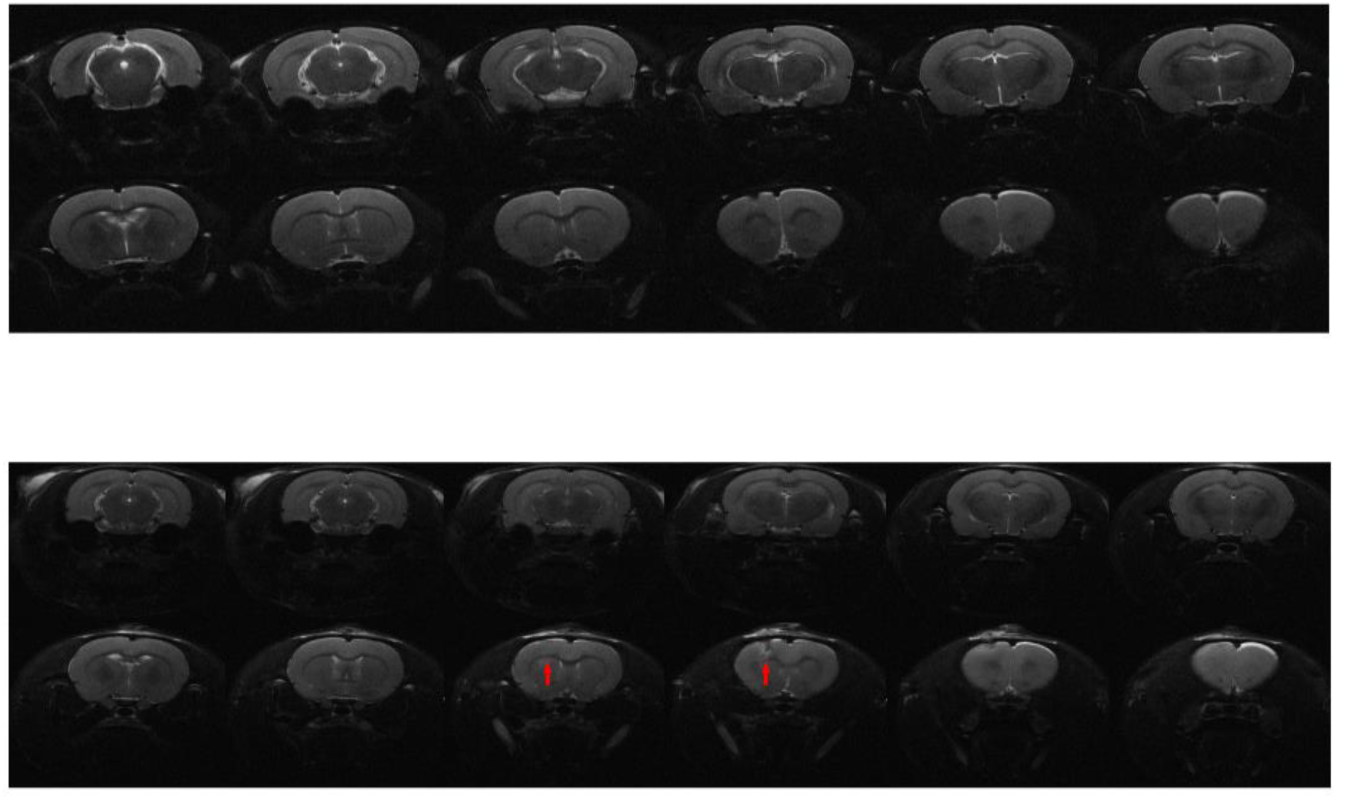
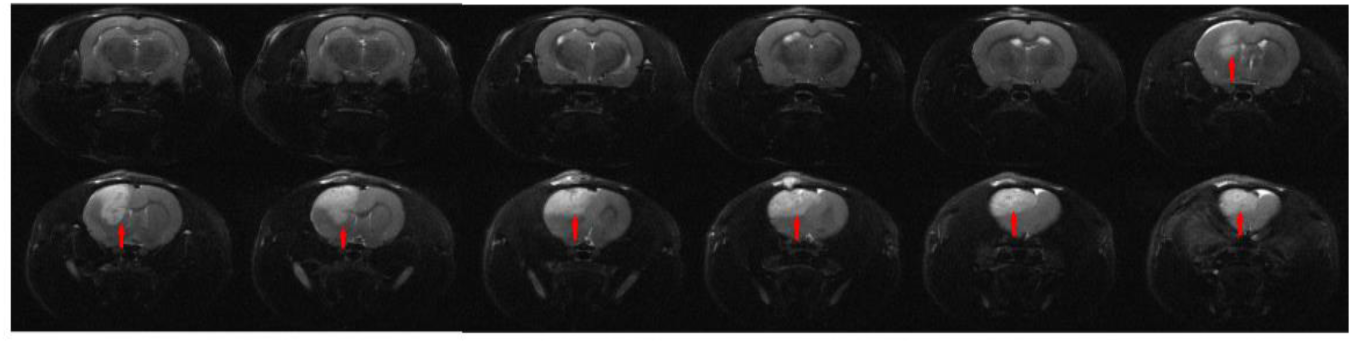
MRI results of the brain tissues after an injection with tumor cells (A) MRI results of the brain tissues on the 6th day. (B) MRI results of the brain tissues on the 10th day. The red arrow indicates the cancer tissues. (C) MRI results of the brain tissues on the 13th day. Red arrows indicate the cancer tissues and the shifted midline.

### Histopathology

Rats were sacrificed and the brain tissue separated. H&E staining is shown. Compared to normal tissues, there is tumor tissue in the right caudate nucleus on the 13th day (Fig. 4A). The tumor tissue in GBM rats exhibited invasive growth with an ill-defined margin (Fig. 4B). Nuclear condensation, fragmentation and pathologic mitosis are observed in the tumor tissue (Fig. 4C).

**Fig 4.**
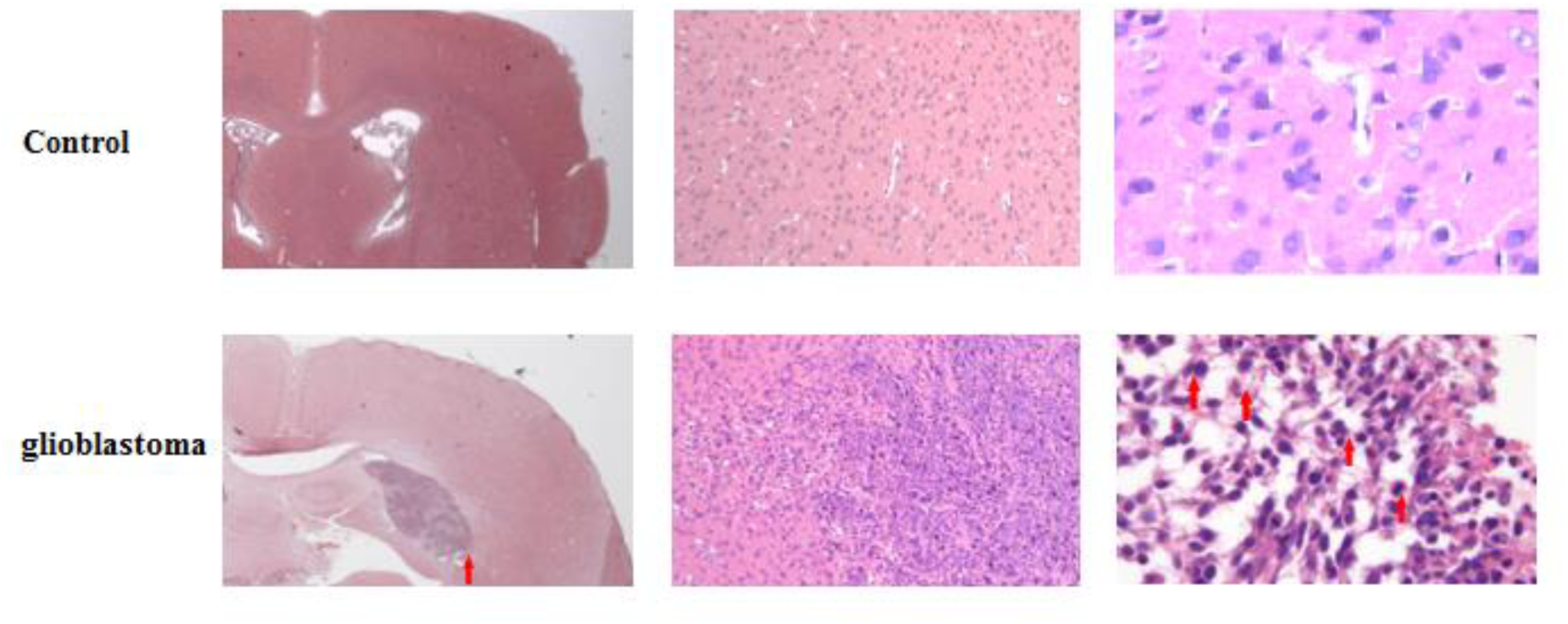
H&E staining of brain tissues on the 13th day after an injection with rat glioma C6 cells. (A) H&E staining (7.5×). The red arrow indicates cancer tissues. (B) H&E staining (100×). (C) H&E staining (400×). The red arrow indicates tumor mitotic figure.

### Differential urinary proteins

The urinary proteins were analyzed using Thermo Orbitrap Fusion Lumos. The urinary samples of three rats were analyzed three times to provide technical replicates (Table S1). In total, 778 proteins were identified (Table S2), and 124 differential proteins were selected according to the following criteria: 1) proteins with at least two unique peptides were allowed, 2) fold change ≥2; and 3) p value < 0.05. All differential proteins were significantly changed in the three rats’ urine.

Among the differential proteins, 109 have corresponding human orthologs, out of which 78 were down-regulated more than 2-fold and 31 were up-regulated more than 2-fold (Table 1). Overall, 56 significantly changed proteins were selected in the 2nd day group (D2), 65 proteins in the D6 group, 61 proteins in the D10 group and 27 proteins in the D13 group, respectively. Among them, sixty-five differential proteins were repeatedly identified in different groups (Fig. S1).

**Table 1.** Number of up-regulated and down-regulated differential proteins in different GBM groups.

| Group | Upregulation | Downregulation | Total |
| --- | --- | --- | --- |
| D2 | 20 | 36 | 778 |
| Percentage | 2.57 | 4.62 | 100 |
| D6 | 4 | 61 | 778 |
| Percentage | 0.51 | 7.84 | 100 |
| D10 | 8 | 53 | 778 |
| **Percentage** | 1.02 | 6.81 | 100 |
| **D13** | 6 | 21 | 778 |
| **Percentage** | 0.77 | 2.69 | 100 |
| **Total (differential proteins)** | 31 | 78 | 109 |

On the 2nd and 6th day, GBM rats showed 90 differential urinary proteins with human orthologs. Among all differential proteins, 27 proteins that had been identified in the CSF, blood or brain tissue were reported to be associated with GBM (Table 2).

**Table 2.**
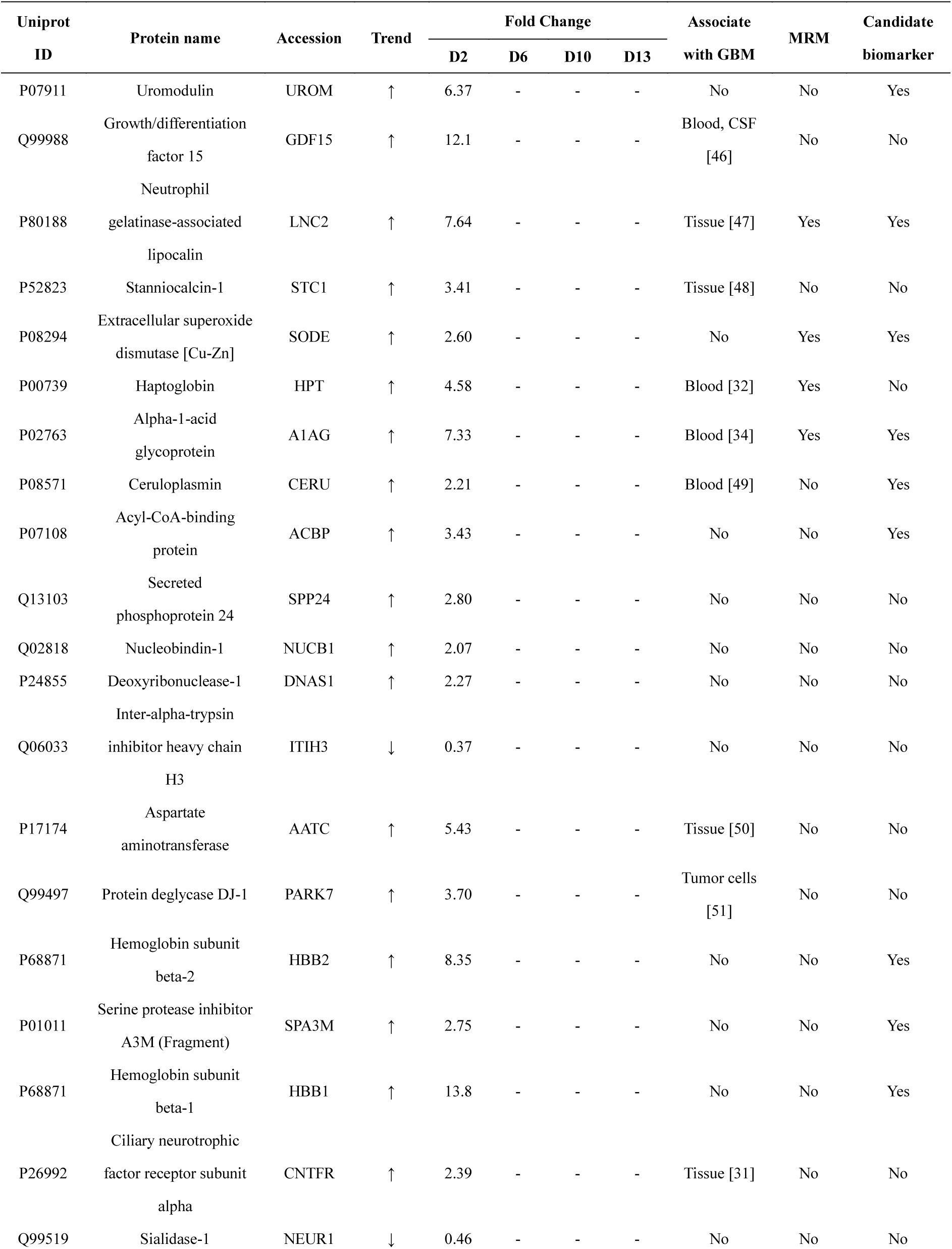
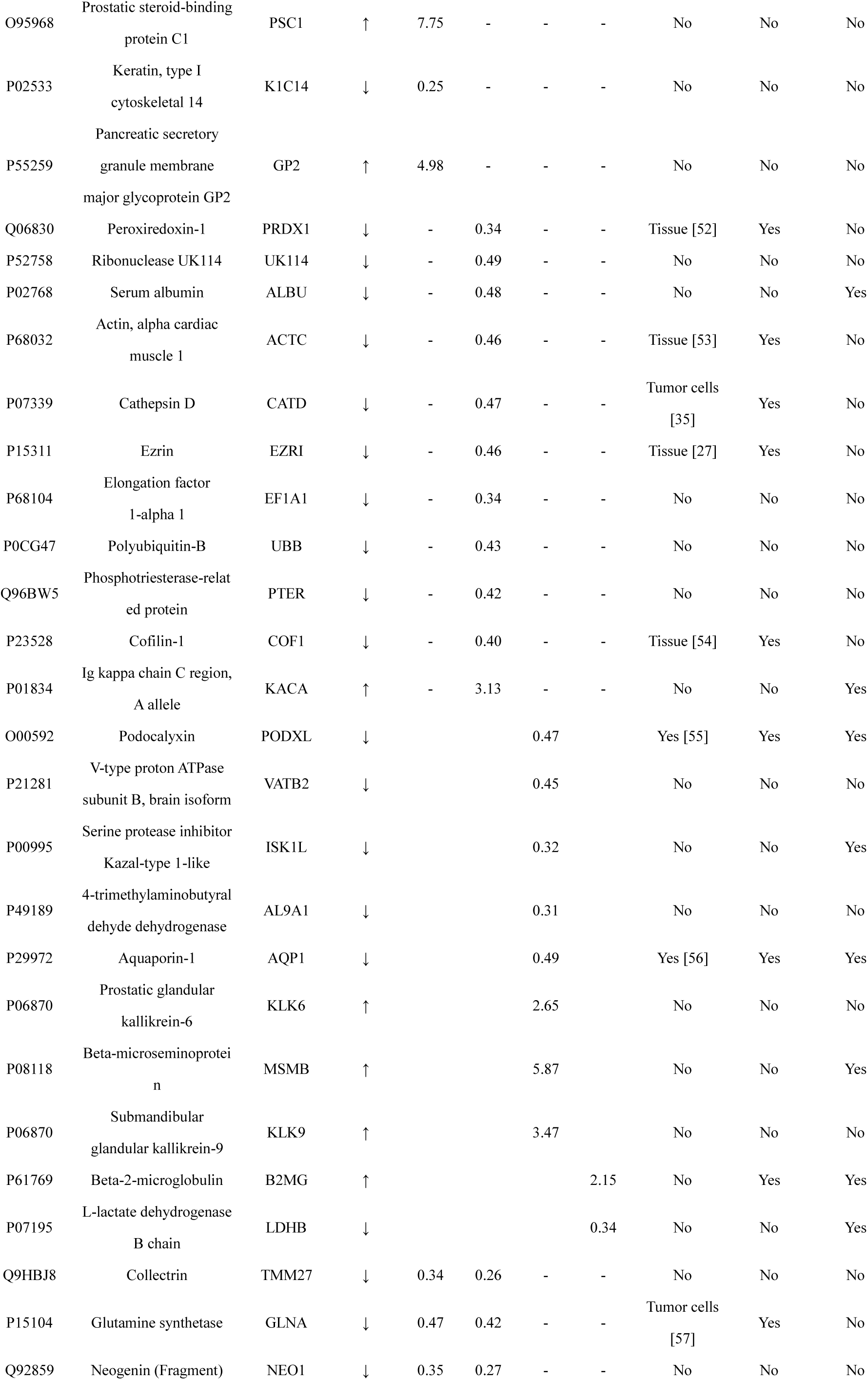
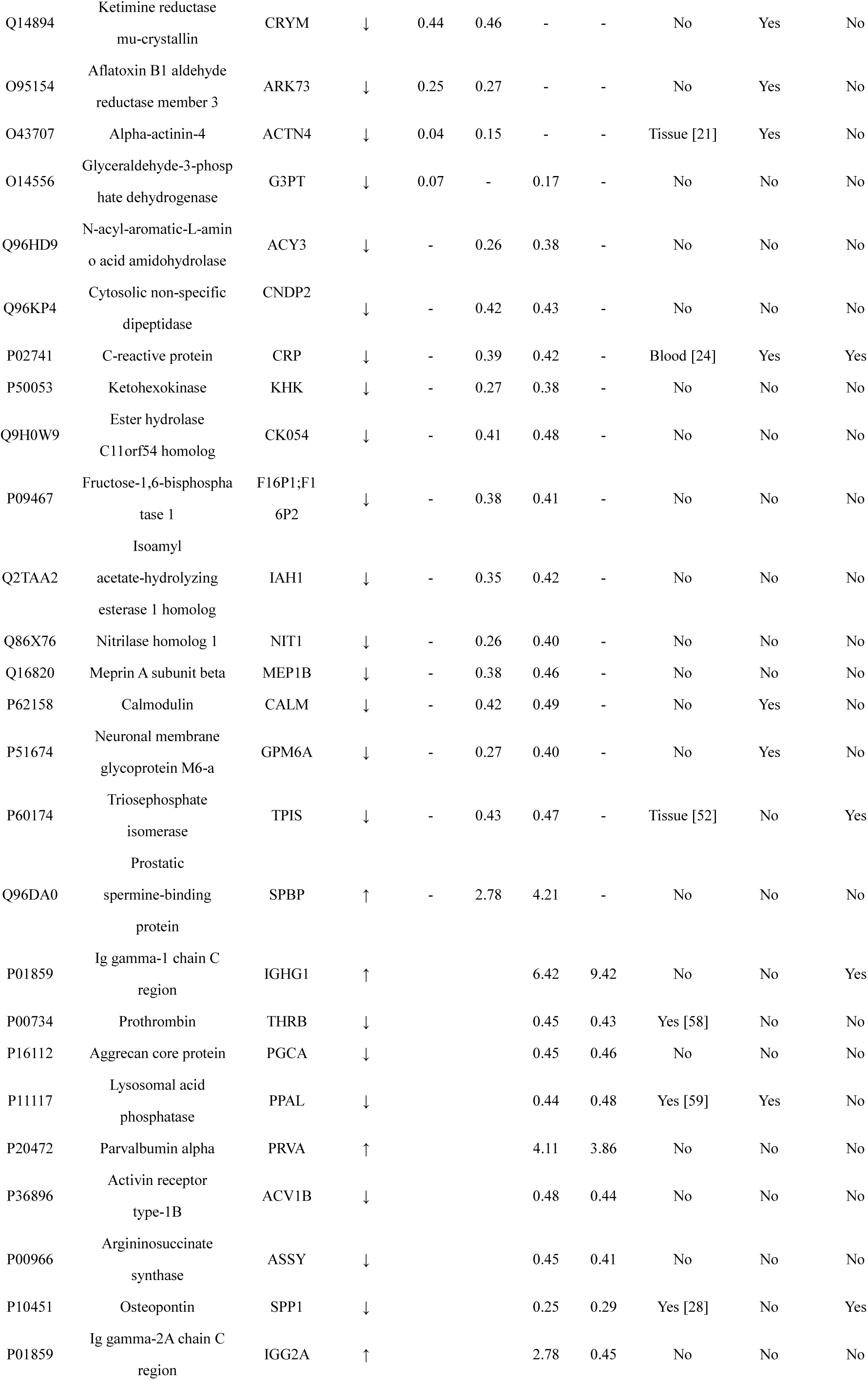
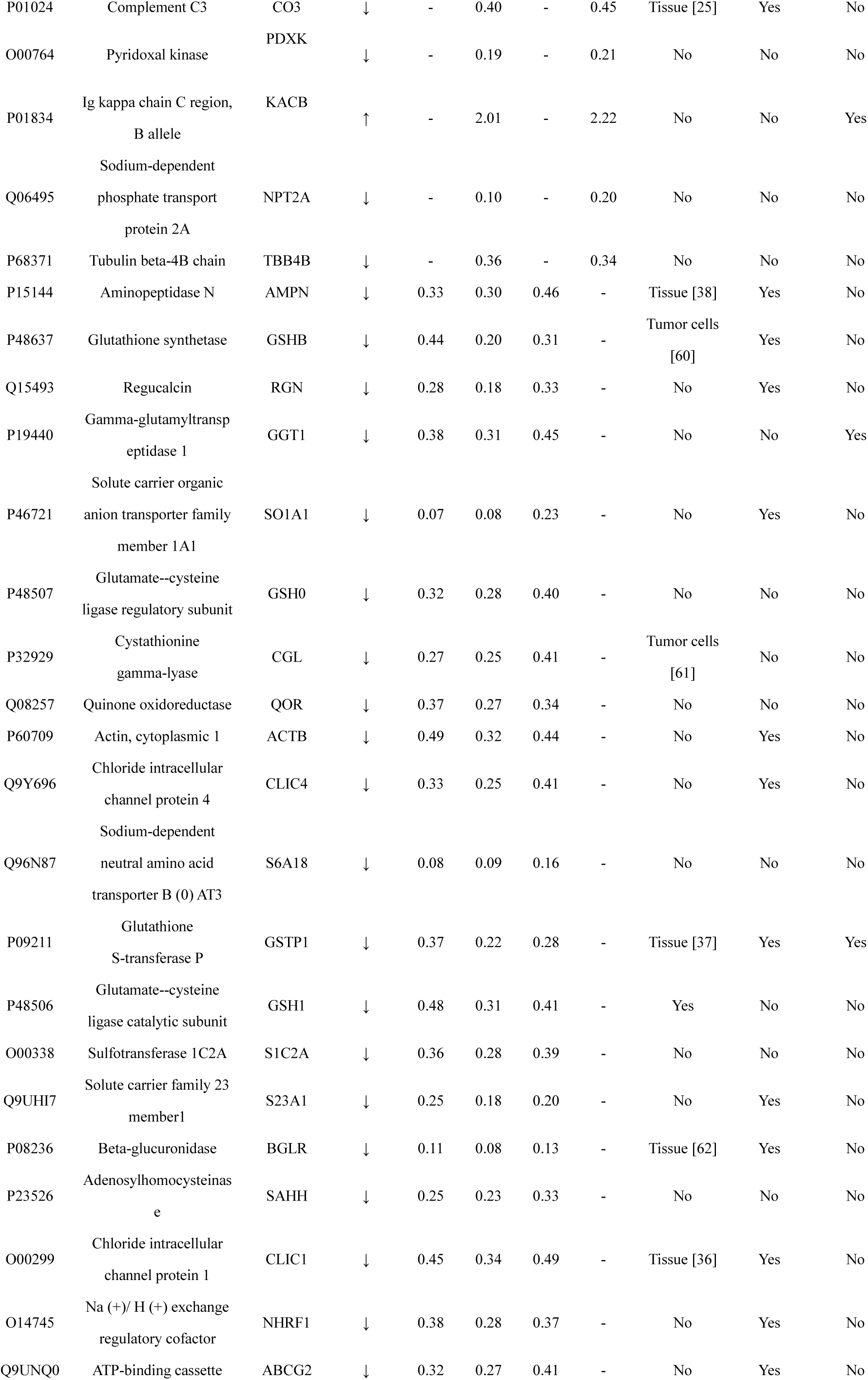
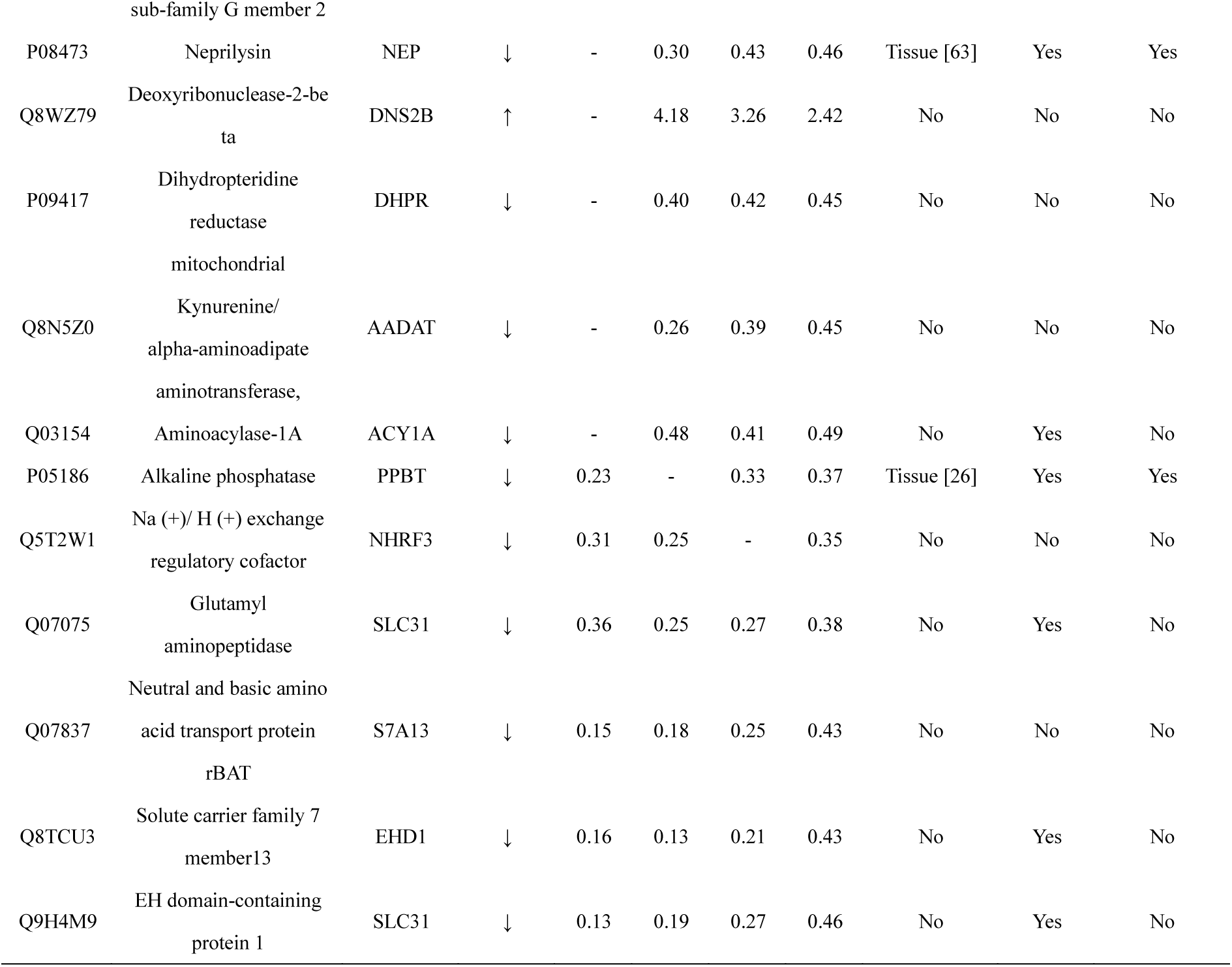
Details of urinary differential proteins identified on the 2nd and 6th day groups.

### MRM analysis

Thirty-nine differential proteins, which were changed in all three rats, were selected for further validation in four additional individual urine samples using MRM. To obtain more confidence, 4-5 transitions were used for each peptide (Table S3). For MRM analysis, the technical variation in the triplicate runs was first calculated and 92.58% of proteins with CV < 0.3 (Table S3), indicating that the technical variation of MRM analysis was ideal.

For the differential proteins, 29 were down-regulated more than 2-fold in the MRM validation results. Glutamyl aminopeptidase was down-regulated in D2, D6, D10 and D13 groups, beta-glucuronidase was also significantly decreased in D2, D6 and D10 groups, which were consistent with MS/MS results (Fig 5). And 9 differential proteins were up-regulated more than 2-fold. Neutrophil gelatinase-associated lipocalin (NGAL) showed a significant elevation in D2 group, beta-2-microglobulin was also significantly increased in D13 group, which confirms the results of our proteome analysis. However, the MRM result of some differential proteins, such as growth and differentiation factor-15 (GDF15) and stanniocalcin-1 which had been reported to be associated with GBM, did not showed significant difference between control and D2 group, which may be due to its lower relative abundance, which requires larger samples and deeper identification for validation.

**Fig 5.**
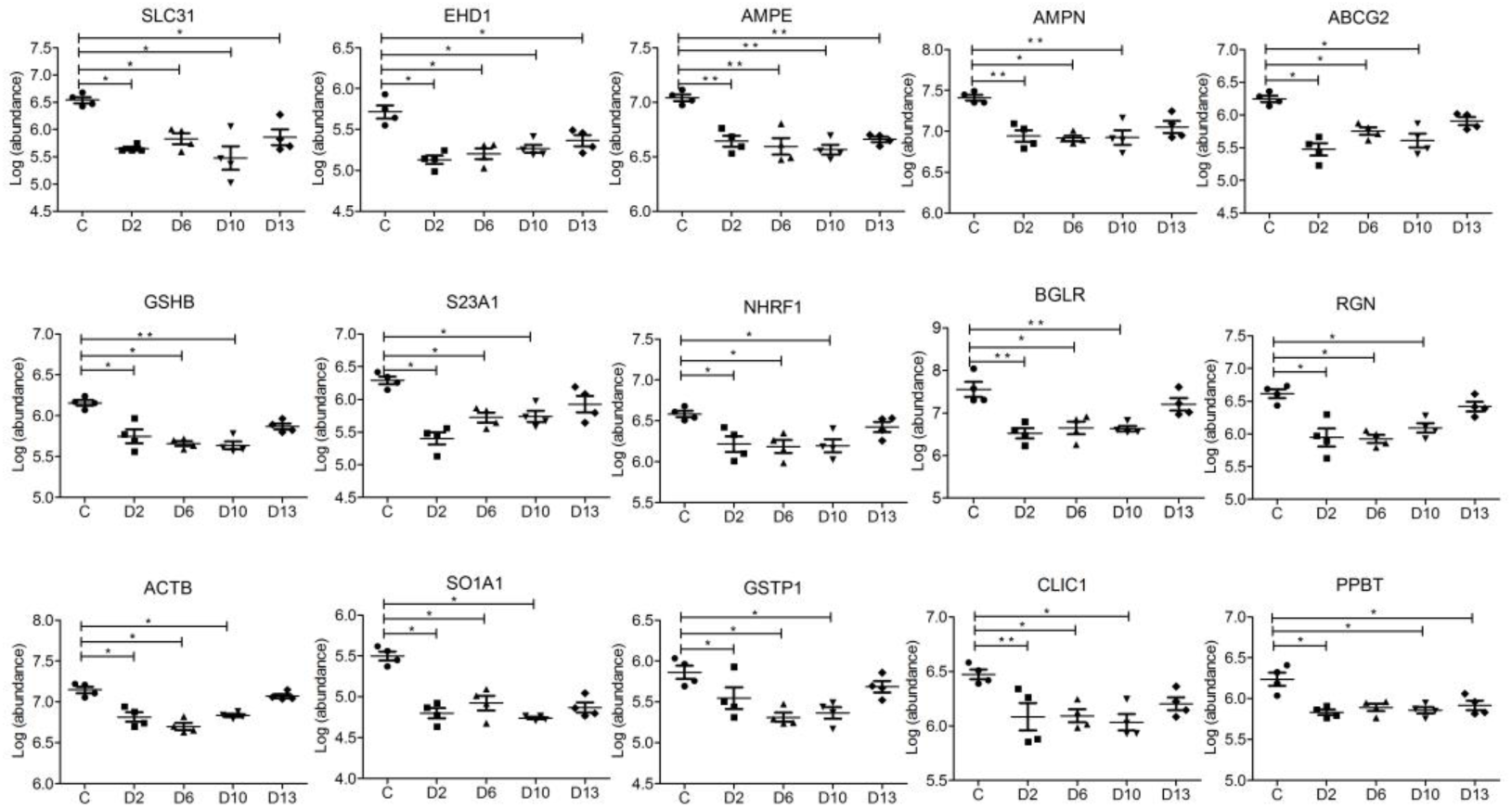
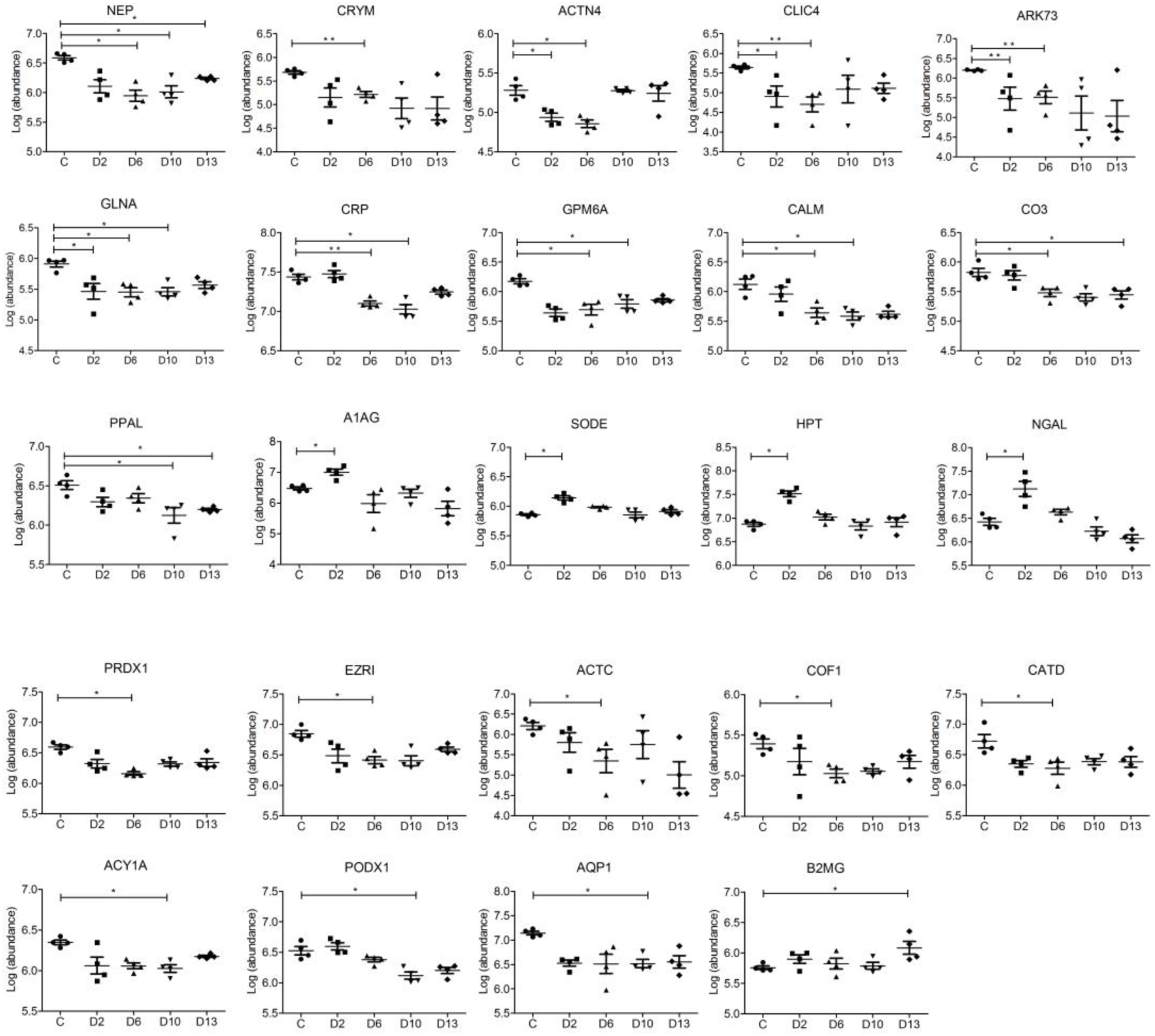
MRM results of candidate urine biomarkers of GBM. Thirty-four proteins shared a decreasing trend in relative abundance, five proteins shared an increasing trend. The x-axis represents different groups; the y-axis refers to the log transformation of the normalized abundance identified by skyline software.

### Gene ontology and Ingenuity Pathway Analysis analyses of differential proteins

The differential proteins were analyzed by the PANTHER classification system. In the biological process category (Fig 6A), the percentages of multicellular organismal processes and metabolic processes were overrepresented, whereas the biological adhesion and response to stimulus were underrepresented in astrocytoma group compared with the whole genome data. In the molecular category (Fig 6B), the percentage of catalytic activity was much higher, while the percentages of receptor and signal transducer activity were much lower in the astrocytoma group in relation to the whole genome data. In the cellular component category (Fig 6C), the membrane was overrepresented in D6/D10 and D13 groups, whereas extracellular region proteins were underrepresented, indicating that the membrane proteins play a role in the pathologic process of astrocytoma.

**Fig 6.**
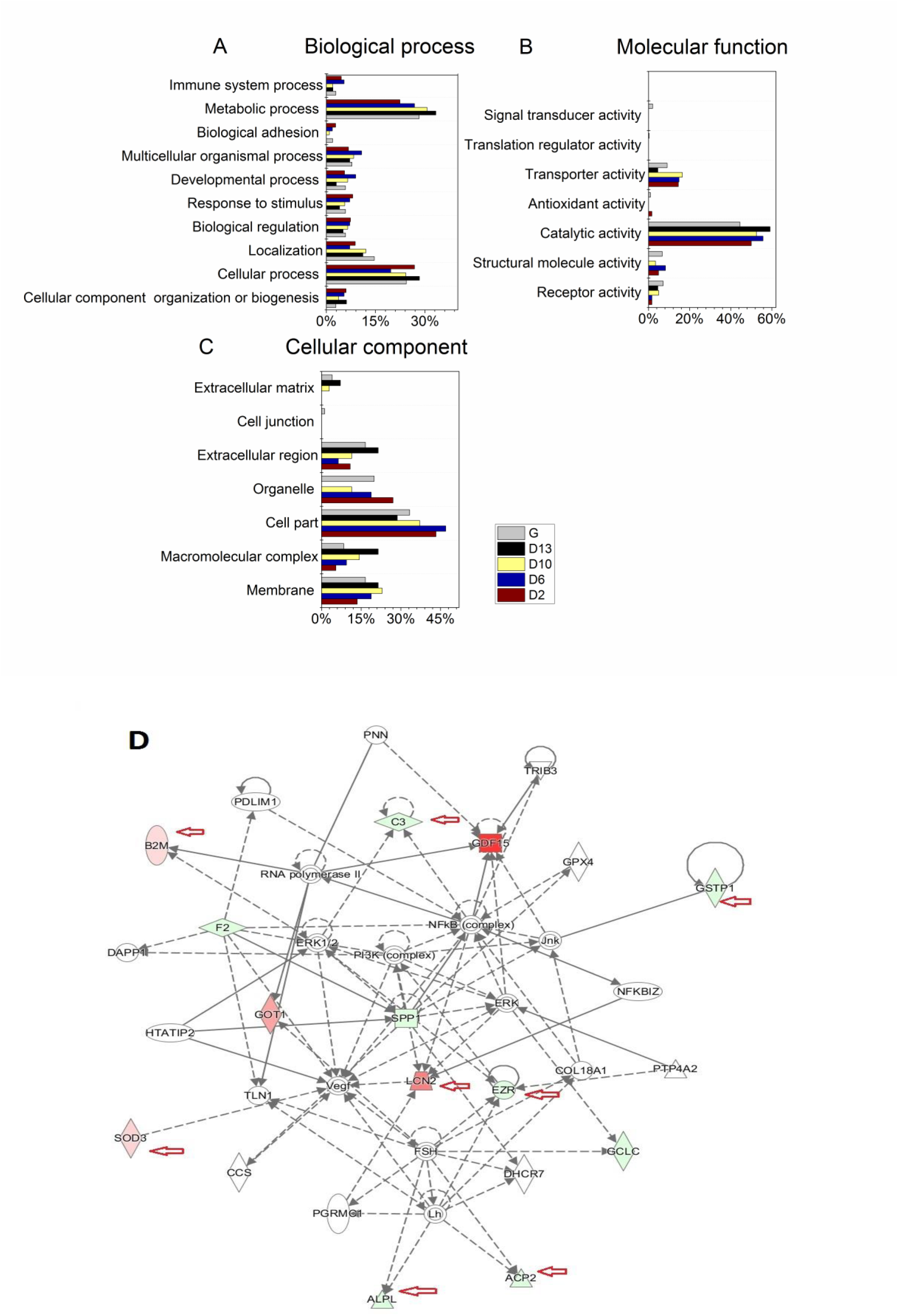
Panther and IPA network analysis of differential proteins. (A) The molecular function analysis between the control and GBM groups; (B) The biological process analysis between the control and GBM groups. (C) The cellular component analysis between the control and GBM groups.(D) Death and Survival networks from IPA analysis. Proteins in red were up-regulated in GBM compared with control rats, and proteins in green were down-regulated in GBM. Proteins pointed by the red arrow were validated using MRM.

Canonical signaling pathways were analyzed to better understand the biological features of glioma. Alpha-actinin-4 (ACTN4), which was involved in VEGF signaling, decreased in the urine of three rats, which is consistent with a previous study’s finding that ACTN4 plays a role in the development and progression of glioma [21]. The glutathione synthetase, which is involved in glutathione biosynthesis, decreased in urine. VEGF signaling and glutathione biosynthesis were enriched in the D2, D6 and D10 groups, and these results were consistent with previous work [22, 23], indicating that our proteomic data may reflect the information of glioma. In the acute phase response signaling, acute-phase proteins, such as complement C3 and C-reactive protein, which may reflect the inflammation and activation of the innate immune system during the course of glioma [24, 25].

The network analysis showed that the cellular death and survival networks were mostly affected (Fig 6D). Osteopontin (SPP1), which is primarily involved in immune responses, tissue remodeling and biomineralization, is a core component of these networks. NGAL (LCN2), extracellular superoxide dismutase (SOD3) and Beta-2-microglobulin (B2M) were up-regulated in these networks. Complement C3 (C3), Glutathione S-transferase P (GSTP1), alkaline phosphatase (ALPL), lysosomal acid phosphatase (ACP2) and ezrin (EZR) were down-regulated in relation to the control group and have been reported to play roles in GBM [26–28].

## Discussion

NGAL complex activity was elevated in brain tumors and somay serve as a molecular marker for brain tumors [29]. NGAL protein may also be involved in glioma drug resistance and clinical prognosis [30]. Ciliary neurotrophic factor receptor subunit alpha, which participates in the formation of tumor-initiating cells in gliomas, is a marker that correlates with histological grade [31]. Serum acute phase reactant proteins were correlated with GBM in relation to their prognosis. Haptoglobin, previously reported to be involved in infection, tumor growth and migration, was identified as a GBM-specific serum marker [32, 33]. Alpha 1-acid glycoprotein, another serum acute phase reactant protein, was also identified in the serum of patient’s with GBM [34]. The lysosomal marker cathepsin D (CATD), which was identified on the 6th day, was frequently overexpressed in glioblastomas [35]. The above differential proteins identified during the early stage may help in providing clues for the early diagnosis of the disease.

Some differential proteins were continuously identified in the D2, D6, D10 or D13 groups. Chloride intracellular channel protein 1 expression presented a correlation with glioma, as higher expression levels were observed in tumor tissues [36]. GSTP1 was reported to show higher expression levels in meningeal tumors and GBM [37]. And a significant decrease in aminopeptidase N (AMPN) occurred concomitantly with tumor growth in glioma tissue [38]. These differential proteins continuously identified in the D2, D6, D10 or D13 groups may help in monitoring the development of the disease.

Most of the differentially expressed proteins that had not been reported to be associated with GBM in other studies may also play an important role and serve as new markers during the progression of this disease. For example, secreted phosphoprotein 24, a bone matrix protein that can suppress pancreatic cancer growth [39] and lung cancer [40], may also be potential urinary markers of GBM.

Several differential proteins may appear in many different brain diseases. For example, AMPN was increased in the urine of both GBM rats and obstructive nephropathy rat [41]; CATD was decreased in the urine of GBM rats but increased in the urine of bladder cancer rats [42], which suggests that these diseases may share similar pathological processes [43]. Thus, it may be difficult to provide an accurate diagnosis using a single biomarker; a panel or collection of urinary proteins may be faster and more sensitive [44]. In this study, one hundred and nine differential proteins with human orthologs could be evaluated together to reflect the progression of GBM.

Compared to the urinary biomarker panel of another cancer, ovarian carcinoma, the protein changes in this study were substantially different. Distinct differential proteins were present in GBM and ovarian carcinoma [45]. The expression profile of differentially expressed proteins in different cancers illustrates that specific pathological conditions have their own biomarker combinations, and different diseases can be differentiated using different biomarker combinations.

In summary, clinical manifestation may provide clues to astrocytoma on the 13th day after tumor cell implantation, while the enhancement lesion appeared in MRI on the 10th day. However, the urinary differential proteins of three rats were changed with the development of the disease and can provide valuable clues by the 2nd and 6th day. Among all the differential proteins in the early stage, 18 proteins were reported to be associated with GBM, suggesting that a panel or collection of urinary differential proteins in urine is a better choice for the early diagnosis of brain diseases (Fig 7). In future studies, we hope this work will help to identify astrocytoma biomarkers with clinical utility and will help to understand the role of urine as a better source in biomarker discovery, especially in brain diseases.

**Fig 7.**
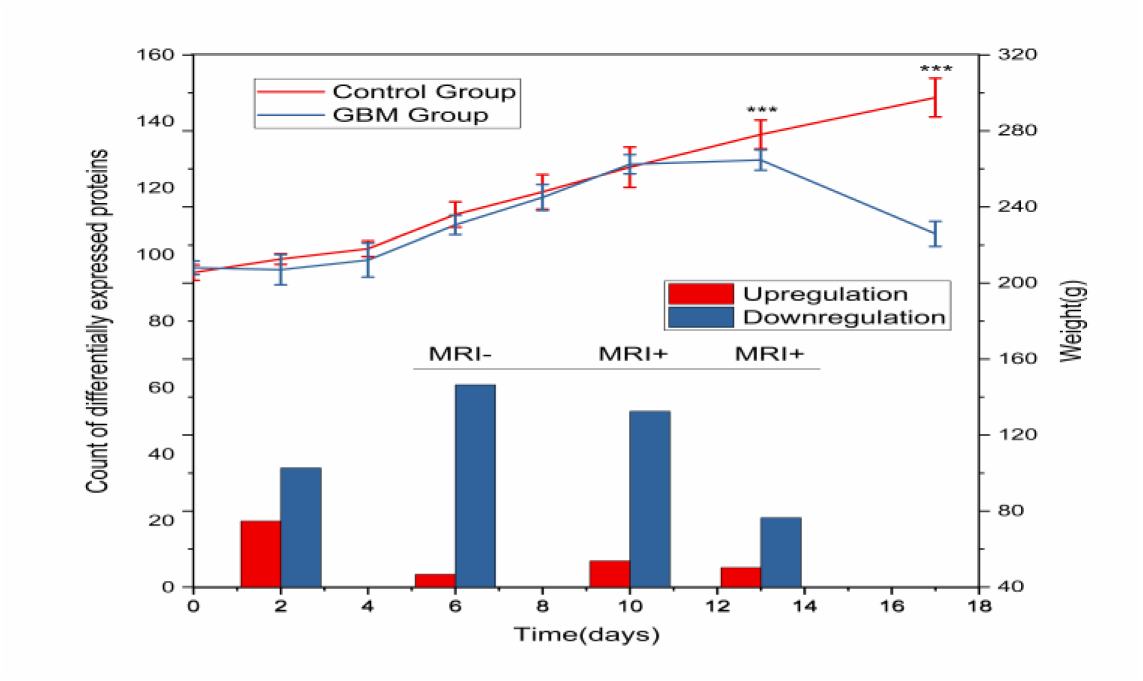
A panel or collection of differential urinary proteins can provide clues for the early diagnose of GBM. Clinical manifestation may provide clues on the 13th day after tumor cell implantation, while the enhancement lesion appeared in MRI on the 10th day. However, the urinary differential proteins of three rats can provide valuable clues by the 2nd and 6th day.

## Acknowledgements

This work was supported by the National Key Research and Development Program of China (2016 YFC 1306300); the National Basic Research Program of China (2013CB530850); Beijing Natural Science Foundation (7173264, 7172076) and funds from Beijing Normal University (11100704, 10300-310421102).

---

**Fig S1.**
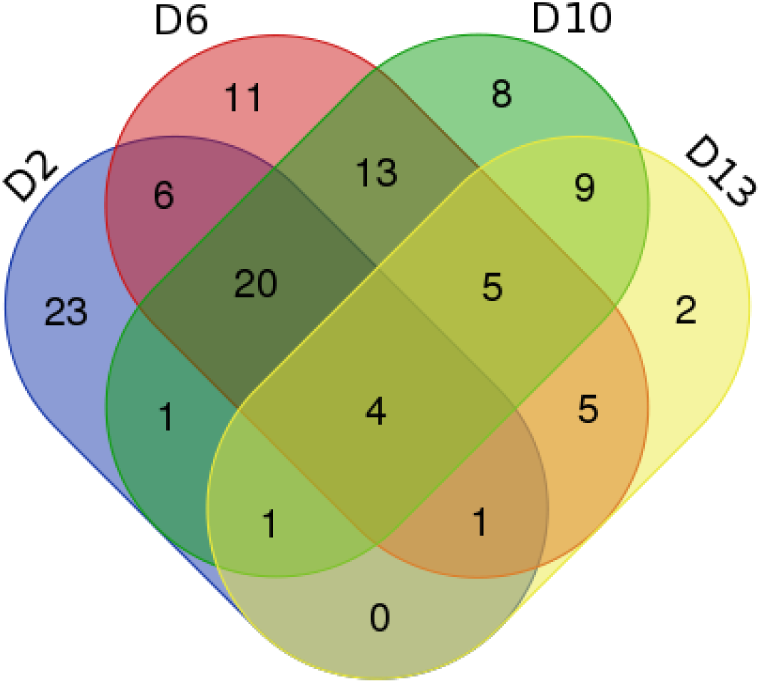
The differential proteins identified in GBM rats. Compared with control group, four proteins were identified in all four GBM groups; twenty-seven proteins were identified in three GBM groups; thirty-four proteins were identified in two GBM groups.

